# Electrocutaneous stimulation to the face inhibits motor evoked potentials in the hand: face-hand interactions revealed by afferent inhibition

**DOI:** 10.1101/719708

**Authors:** Bia L. Ramalho, Julien Moly, Estelle Raffin, Sylvain Harquel, Alessandro Farnè, Karen T. Reilly

**Author notes:** Laboratory of Neurobiology II, Institute of Biophysics Carlos Chagas Filho, Federal University of Rio de Janeiro, Rio de Janeiro, Brazil. Corresponding authors, (BR) and (KR).

## Abstract

Reorganization of the sensorimotor cortex following amputation and other interventions has revealed large-scale plastic changes between the hand and face representations. To investigate whether hand-face interactions are also present in the normal state of the system we measured sensorimotor interactions between these two areas using an afferent inhibition transcranial magnetic stimulation (TMS) protocol in which the TMS motor evoked potential (MEP) is inhibited when it is preceded by an afferent stimulus. We hypothesized that if hand-face interactions exist in the normal state of the system then stimulation of the face would inhibit hand MEPs. In two separate experiments we delivered an electrocutaneous stimulus to either the right upper lip (Experiment 1) or right cheek (Experiment 2) and recorded muscular activity from the right first dorsal interosseous (FDI). Both lip and cheek stimulation inhibited FDI MEPs. To investigate the specificity of this effect we conducted two additional experiments in which cutaneous stimulation was applied to either the right forearm (Experiment 3) or right arm (Experiment 4). Neither forearm nor arm stimulation inhibited FDI MEPs. These data provide the first evidence for face-to-hand afferent inhibition and we suggest that the mechanisms underlying these sensorimotor interactions could contribute to face/hand interactions observed following sensorimotor reorganisation.

## Introduction

The sensory and motor cortices contain somatotopic maps of body areas and muscles. The medio-lateral organization of these maps is such that the lower limb is represented medially followed by the trunk, upper limb, and most laterally the hand and face. The face and hand representations differ from those of other body parts because their proximity within the maps contrasts starkly with their separation in the physical body. Furthermore, the extensive area devoted to processing stimuli or controlling musculature from these two body parts is disproportionate to their physical size [1,2].

The physical proximity of the face and hand within somatosensory and motor maps is thought to underlie the plasticity that has been documented following a reduction of sensorimotor inputs from the hand. For example, many studies have found that amputation or temporary nerve block of the hand induces an enlargement and shift of the face’s sensorimotor representation [3–7]. This plasticity is paralleled by behavioural changes like referred sensations in the phantom hand following face stimulation in hand amputees [5,8], phantom limb pain (Karl et al. 2001), face-hand somatosensory extinction after hand allograft [9], or improved two-point discrimination in the upper lip after anaesthesia of hand nerves (median and radial nerves) [4]. While this plasticity has typically been interpreted as resulting from the reduction of inputs, an alternative hypothesis posits that it is driven instead by increased input from other body parts, for example by overuse of the non-amputated hand following upper-limb amputation [10].

In line with the idea that plasticity can also be induced by increased input, our group has demonstrated in healthy subjects that plasticity between the face and hand is not restricted to instances of reduced input, but can also occur after an *increase* in somatosensory inputs. Using repetitive somatosensory stimulation at the right index finger tip Muret et al. (2014) found improved two-point discrimination at the stimulated fingertip but also on the cheek and upper lip. This “transfer” of behavioural improvement was accompanied by alterations in both hand and face representations in the sensory cortex [12].

Face-hand interactions at the somatosensory level are not restricted to situations involving plasticity, but also occur when the system is in its “normal” state. For example, Tanosaki et al., (2003) found that somatosensory evoked magnetic fields induced by electrical stimulation of the thumb were altered when there was concurrent tactile stimulation of the upper face. These findings raise the question of whether face-hand *sensorimotor* interactions might also exist in physiological conditions, in which case they could represent one of the mechanisms underlying both large-scale amputation-induced plasticity as well as temporary, experimentally-induced plasticity, like that observed after repetitive somatosensory stimulation or anaesthesia.

We assessed Short-latency Afferent Inhibition (SAI) to test for the existence of face-hand sensorimotor interactions under normal physiological conditions. SAI is the reduction in the amplitude of a muscle response (evoked by TMS of the motor cortex) when motor cortex stimulation is preceded by an afferent stimulus [14–25]. This protocol can provide information about latent sensorimotor interactions between body parts [26–29], and has been widely used to examine sensory interactions *within* the same body part. For example, hand muscle responses are strongly inhibited following stimulation of hand nerves or the skin on the fingertip [14,15,24,25,27,28,16–23], especially when the stimulus is given close to the target muscle [30], and topographic information can be preserved in the sensory-to-motor inhibitory pattern [31]. Similarly, stimulation of the shoulder area inhibits responses in the shoulder muscle infraspinatus [32], and stimulation of the dorsal surface of the foot inhibits responses in the leg muscle tibialis anterior [29]. Afferent inhibition can also occur when the muscle of interest and the sensory stimulation site are within the same body part but anatomically separate. For example, stimulation of the index fingertip inhibits various muscles of the arm and forearm on the same side as the fingertip stimulation [28], and can also inhibit hand muscles on the opposite side of the body [29].

To date, there is no evidence for the existence of SAI between body parts, although the only combinations examined have been the lower and upper limbs [28]. Here we investigated if and when SAI exists between the face and the hand, as it is known that these anatomically distant body parts have strong interactions both under normal physiological conditions and following a plasticity-inducing manipulation. SAI is typically considered to occur at a latency related to the delay of arrival of the afferent information at the motor cortex. For example, following fingertip stimulation, maximal inhibition is observed between 25 and 35 ms. This assumption is based upon within-body part SAI, however, there are currently no data indicating whether a similar rule applies for SAI between body parts. Furthermore, the results of the only study that investigated SAI within the face suggest that face stimulation might not follow this same rule, as there was some evidence of face-face SAI at 30ms but none at shorter (expected) ISIs [22]. Thus, the aim of the experiments presented here was to 1) establish **if and when** afferent inhibition exists between two anatomically separate body parts: the face and the hand; and 2) whether this between body-part inhibition is specific to the face and the hand or is also present between the forearm or upper-arm and the hand.

## Materials and Methods

### Participants

Forty-four healthy right-handed volunteers were included in four separate experiments. It is important to note that each experiment was independent of the others, as the aim of this study was to investigate if and when SAI exists between a given body part and the right FDI, not to compare the amount or latency of SAI between the four stimulated sites. Fourteen individuals participated in Experiment 1 (mean age of 22.7 ± 7.1 years, 5 males), 12 in Experiment 2 (mean age of 23.7 ± 6.7 years, 3 males), 15 in Experiment 3 (mean age of 25.5 ± 6.4 years, 2 males) and 13 in Experiment 4 (mean age of 24.8 ± 3.9 years, 3 males). Four subjects participated in two experiments (1 & 2 (n=2); 1 & 3 (n=1); 3 & 4 (n=1)) and 3 subjects participated in 3 experiments (1, 2 & 3 (n=1); 1, 3 & 4 (n=1); 2, 3 & 4 (n=1)). All participants gave written informed consent. The protocol was approved by the ethical committees of the Grenoble University Hospital (ID RCB: 2016-A01668-43) and the *Comité de protection des personnes* (CPP) SUD EST IV (ID RCB: 2010-A01180-39) and conformed to the ethical aspects of the Declaration of Helsinki.

### General experimental procedures

In each of the four experiments participants were comfortably seated with their arm resting on an armrest (elbow flexed at 90°) and a single tactile electrical stimulus was applied prior to a single transcranial magnetic stimulation pulse over the hand area of the left motor cortex. The tactile stimulus was applied to the right upper lip (Experiment 1), right cheek (Experiment 2), right forearm (Experiment 3), or right arm (Experiment 4). In all four experiments electromyographic activity was recorded from the right first dorsal interosseous (FDI) and the inter-stimulus intervals (ISIs) between the electrocutaneous stimulus and the TMS pulse were 15, 25, 35, 45, 55, 65, 75 and 85 ms. In all four experiments 14 trials of each ISI plus 34 TMS-only trials were presented in a random order with an inter-trial interval between 5 and 8 seconds. Every 24 trials the experiment was paused to give a short break to the participant. Experiments 1 and 2 were conducted in the IMPACT team (Lyon, France) and Experiments 3 and 4 in the IRMaGe MRI and Neurophysiology facilities (Grenoble, France).

### Electrical Stimulation

Single pulse electrocutaneous stimuli (square wave, 200 μs) were delivered via a constant current stimulator (DS7A, Digitimer Ltd, UK) using bipolar adhesive electrodes (Neuroline 700, Ambu, Copenhagen, Denmark) placed on the face (Experiments 1 & 2) or the upper limb (Experiments 3 & 4). The sensory perception threshold (SPT) for each stimulation site was determined as the minimum stimulation intensity at which the subject reported feeling the stimulation on 2 out of 3 trials. Sensory afferent inhibition protocols always use non-painful stimuli and typically use intensities between 2 and 3 times SPT [14,26,28,31,33]. Tamburin et al. (2001) showed that stimulation applied to the tip of the little finger at 3xSPT produced inhibition in abductor digiti minimi comparable to that recorded at 5xSPT, and Bikmullina et al. (2009) showed that when stimulation was applied to the index finger inhibition in arm and forearm muscles was greater at 3xSPT than at 1x or 2x. The electrocutaneous stimulus intensities used in each experiment are shown in Table 1 and the Kruskal-Wallis test by ranks revealed no difference between the absolute stimulus intensity used in each of the four experiments (p= 0.15).

**Table 1.** Average electrocutaneous and TMS stimulus intensities used in each of the four experiments, plus average TMS-only amplitude of FDI MEPs (mean ± SEM).

| Exp. | ES location | ES intensity<br>(mA) | rMT<br>(%MSO) | 1mV intensity<br>(%MSO) | TMS-only<br>(mV) |
| --- | --- | --- | --- | --- | --- |
| 1 | Upper Lip | 4.1 ± 0.5 | 40 ± 1.0 | 44 ± 2.0 | 0.9 ± 0.1 |
| 2 | Cheek | 5.0 ± 0.4 | 39 ± 0.1 | 43 ± 0.2 | 1.0 ± 0.2 |
| 3 | Forearm | 4.6 ± 0.2 | 45 ± 3.6 | 57 ± 4.6 | 1.1 ± 0.1 |
| 4 | Arm | 4.1 ± 0.3 | 47 ± 2.8 | 59 ± 3.6 | 1.1 ± 0.1 |

ES: Electrocutaneous Stimulation; rMT: resting Motor Threshold; MSO: Maximum Stimulator Output; TMS-only: FDI MEP amplitude in the absence of electrocutaneous stimulation.

#### Experiment 1: SAI between the right upper lip and the right FDI

Two electrodes were placed side-by-side horizontally, separated by 1 cm, over the right upper lip with the more medial electrode close to the phitral ridge. Stimulation was delivered at 2xSPT because higher intensities were reported as painful in the majority of subjects.

#### Experiment 2: SAI between the right cheek and the right FDI

Two electrodes were placed 1 cm apart vertically, at the approximate midpoint between the right ear and the right corner of the mouth. As for the upper-lip, stimulation was delivered at 2xSPT because higher intensities were reported as painful in the majority of subjects.

#### Experiment 3: SAI between the right forearm and the right FDI

Electrodes were placed 1 cm apart on the anterolateral face of the forearm in the middle of the proximal third of the forearm on the skin overlying the extensor carpi radialis. Stimulation was delivered at 3xSPT.

#### Experiment 4: SAI between the right arm and the right FDI

Electrodes were placed 1.5 cm apart on the medial face of the arm in the middle of the proximal third of the upper arm on the skin overlying the border between the biceps and the triceps. As for the forearm, the stimulation was delivered at 3xSPT.

### Transcranial magnetic stimulation (TMS) and electromyography (EMG)

TMS was applied over the left motor cortex and EMG activity was recorded from the right FDI via surface electrodes (DE-2.1, Delsys, Massachusetts, USA) placed on the muscle belly. EMG activity was recorded at 2000 Hz, digitized (Power 1401II, Cambridge Electronics Design, Cambridge, UK) and stored on a computer for off-line analysis (Spike 2 or Signal, Cambridge Electronics Design, Cambridge, UK). In Experiments 1 and 2 TMS was applied using a 9 cm figure-of-eight coil and a Magstim 200 stimulator (Magstim, Carmarthenshire, UK). In Experiments 3 and 4 TMS was applied using a 7.5 cm figure of eight coil and a MagPro x100 stimulator (Magventure, Skovlunde, Denmark).

The coil was positioned over the hand area of the primary motor cortex and the optimal point for stimulating FDI was found by stimulating at a slightly suprathreshold intensity and identifying the point with the largest, most stable responses. To enable the experimenter to accurately maintain the coil over the optimal position throughout the experiment this point was recorded in a neuro-navigation system (Brainsight, Rogue Resolutions, Cardiff, UK (Experiments 1 & 2), Localite neuronavigation system, Localite GmbH, Sankt Augustin, Germany (Experiments 3 & 4)). The resting Motor Threshold (rMT) was determined as the minimum stimulator intensity necessary to evoke MEPs of at least 50 μV (peak-to-peak amplitude) on at least 5 out of 10 trials. The TMS pulse intensity used during the experiment was adjusted to produce MEPs in the control condition (TMS-only) with a mean amplitude of approximately 1mV. The average rMT and intensity that produced a MEP of approximately 1mV (both expressed as a percentage of the maximum stimulator output (%MSO)) are shown separately for each experiment in Table 1. A Kruskal-Wallis test revealed no difference between the amplitude of the TMS-only MEPs in each of the four experiments (p= 0.46).

Throughout the experiment, the baseline EMG signal was constantly monitored to ensure that the muscle was completely relaxed. If muscle activity was detected the subject received a verbal instruction to relax the hand. Trials contaminated by muscle contraction in the 500ms before the TMS pulse were excluded from further analyses.

### Statistical analysis

Data from each of the four experiments were analysed separately. Peak-to-peak MEP amplitudes (mV) were measured off-line using custom-written Spike 2 or Signal scripts (Cambridge Electronics Design, Cambridge, UK). Trials were excluded if they were contaminated by muscle contraction or if their amplitudes were greater than or less than 1.96 SDs from the mean of that condition for that subject. On average 16 (± 1.1 SEM) trials were excluded for each subject. The mean MEP amplitude for each condition was then calculated. D’Agostino-Pearson omnibus tests were applied to verify if the data came from an approximately normal distribution. Since the data for some conditions were not normally distributed, a Friedman repeated measures, non-parametric rank test with one factor (ISI) was applied to the raw MEP amplitudes (mV) to compare the mean amplitudes across conditions. Dunn’s Multiple Comparison post-hoc tests comparing the control condition (TMS-only) with each ISI (15 to 85ms) were applied if the factor ISI was significant with a significance level of 0.05. Data were analysed using Prism 5 (GraphPad Software, Inc., California, USA). For each subject, mean MEP amplitude values for each ISI were normalized to the mean of the TMS-only condition and these normalized data were used to graphically represent the results but all analyses were conducted on raw MEP amplitudes.

## Results

### Hand muscle inhibition following electrocutaneous stimulation on the face

#### Experiment 1

Fig 1A shows that electrocutaneous stimulation of the right upper lip inhibited right FDI MEPs by between 20 and 30% at the 45, 55, and 65ms ISIs. A Friedman test on the mean MEP amplitude for each subject in each condition revealed a significant main effect of ISI (χ² (8) = 21.20; p = 0.007). Dunn’s post-hoc tests comparing the mean MEP amplitude at each ISI against the mean TMS-only MEP amplitude revealed that inhibition was significant only at the 45ms ISI.

**Fig 1.**
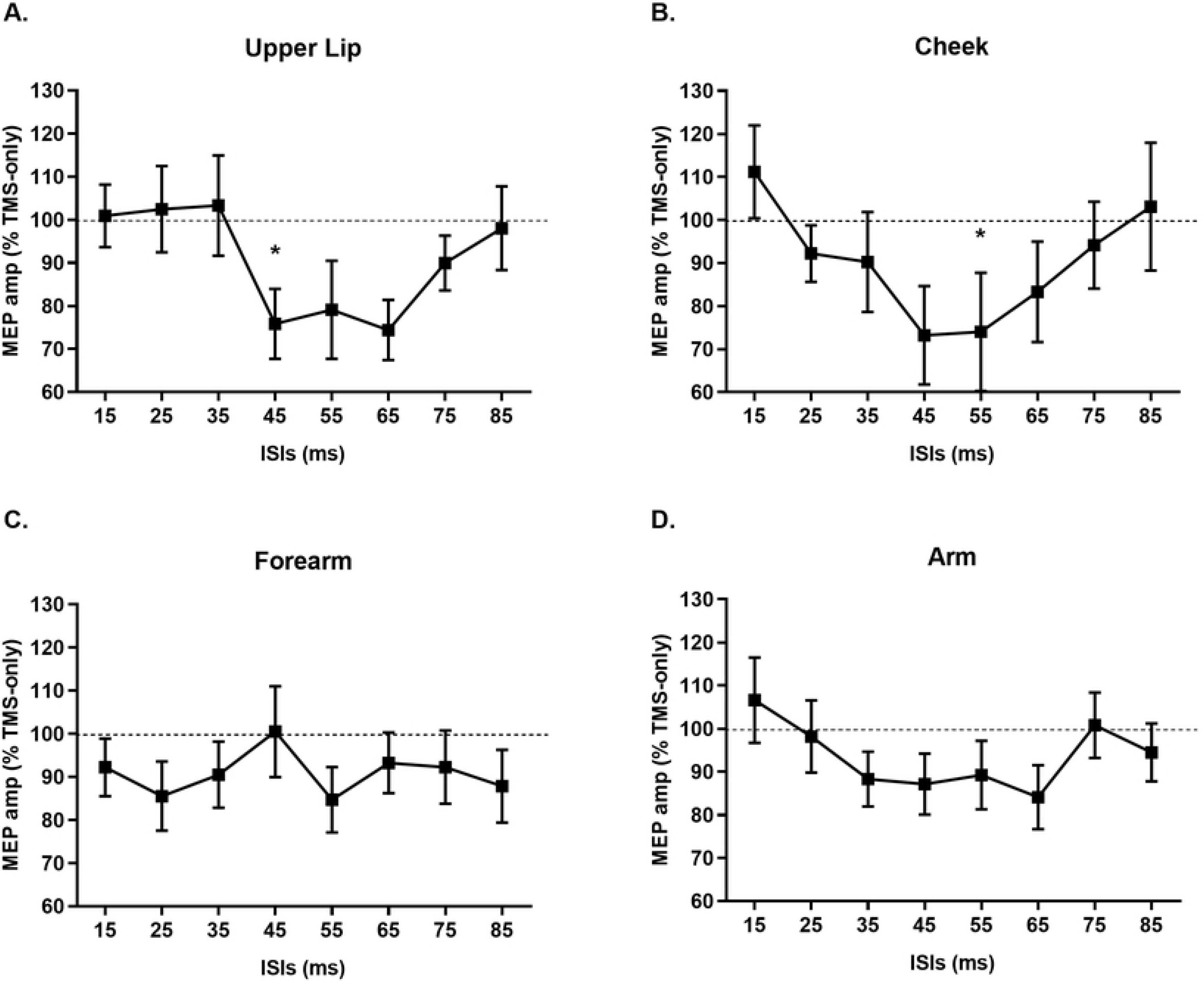
Normalized mean MEP amplitudes in the right FDI after electrocutaneous stimulation of the right upper lip – Experiment 1 (A), right cheek – Experiment 2 (B), right forearm – Experiment 3 (C) and right arm – Experiment 4 (D). Bars represent the standard error of the mean. The black dashed lines represent the TMS-only MEP amplitude. Asterisks represent significant Dunn’s post-hoc tests (p < 0.05) comparing mean TMS-only amplitude with mean amplitude at each ISI. Note that statistical tests were performed on non-normalized data (S1 Appendix).

#### Experiment 2

Fig 1B shows that electrocutaneous stimulation of the right cheek produced a similar pattern and amount of inhibition as lip stimulation (between 20 and 30% at the 45, 55, and 65ms ISIs). A Friedman test on the mean MEP amplitude for each subject in each condition revealed a significant main effect of ISI (χ² (8) = 16.44; p = 0.036). Dunn’s post-hoc tests comparing the mean MEP amplitude at each ISI against the mean TMS-only MEP amplitude revealed that this inhibition was significant only at the 55ms ISI.

### Hand muscle inhibition following electrocutaneous stimulation on the arm

#### Experiment 3

Fig 1C shows that electrocutaneous stimulation of the right forearm inhibited right FDI MEPs by between 10 and 20% at the 25 and 55ms ISIs. A Friedman test on the mean MEP amplitude for each subject in each condition revealed no main effect of ISI (χ² (8) = 8.34; p = 0.401).

#### Experiment 4

Fig 1D shows that electrocutaneous stimulation of the right arm also inhibited FDI MEPs by between 10 and 20% at the 35 to 65ms ISIs, but similar to the forearm, a Friedman test on the mean MEP amplitude for each subject in each condition revealed no main effect of ISI (χ² (8) = 8.96; p = 0.345).

## Discussion

### Face stimulation can inhibit hand MEPs

Face-hand sensorimotor interactions are clearly important for feeding, grooming, non-verbal communication and many other activities of daily life [34]. These interactions exist at a fundamental level in the nervous system in the form of reflexes. For example, the Babkin reflex in neonates occurs when palm pressure evokes mouth opening [35], and the palmomental reflex occurs in adults when thenar eminence stimulation evokes contraction of the mentalis muscle of the chin [36]. Higher-order face-hand interactions have also been documented under situations of plasticity, but evidence for non-reflexive interactions under normal, physiological conditions in the adult is rare. Here, we present the first evidence of sensorimotor afferent inhibitory interactions between the face and the hand. We found that electrocutaneous stimulation of the right upper lip (Experiment 1) and right cheek (Experiment 2) significantly inhibited MEP amplitudes in the right FDI. Interestingly, this between body part SAI appears to be specific to the face and the hand, as despite being anatomically closer to the FDI, forearm (Experiment 3) and arm (Experiment 4) stimulation did not alter the amplitude of FDI MEPs.

These results provide the first evidence for sensorimotor afferent inhibitory interactions between the face and the hand. These findings reinforce the idea that there are privileged interactions between the face and the hand and that such interactions are not limited to reflexes or situations in which the system is perturbed.

The temporal dynamics with which touch on the face inhibited hand muscle responses suggest that face-hand afferent inhibition mechanisms differ from those underlying hand-hand inhibition. For example, most studies examining fingertip or median nerve stimulation show that inhibition of hand muscle MEPs begins just after the arrival of the afferent volley in the primary somatosensory cortex (S1) – at an ISI of approximately 25ms [14,23,27,28]. Many studies even use electroencephalography to measure the latency of this afferent volley and then choose their sensory-TMS ISIs so that the TMS pulse arrives at the time when the sensory information is presumed to have been transferred to the motor cortex i.e. several milliseconds after the arrival of the afferent volley in S1 [20,21,37–40]. This technique is based upon the hypothesis that afferent inhibition results from the activation of direct inhibitory connections from the primary sensory to primary motor cortices. If face-hand afferent inhibition were based upon the same mechanisms as hand-hand afferent inhibition we would have observed it around 15ms, not 45 or 55 ms [41,42]. Interestingly, in a study of face-face afferent inhibition, Pilurzi et al., (2013) found no statistically significant inhibition, but visual inspection of their results (see Fig 5, page 1898) suggests that some inhibition might be present around 30ms – later than would be expected based upon the arrival of the afferent volley in S1 - but similar to the delay we observed for inhibition between the face and the hand. This suggests that the ISIs at which we observed significant face-hand inhibition might not be attributable to the fact that the AI was between two body parts, but might instead be a feature of AI involving face stimulation.

When a somatosensory stimulus arrives in the S1 cortex, it evokes a series of positive (P) and negative (N) deflections. Face stimulation evokes somatosensory evoked potentials (SEPs for EEG) or somatosensory evoked fields (SEFs for MEG) between 15ms (N15) and 65ms (P65) [41–45]. Other deflections are also measurable at longer latencies (70-120ms), and these are thought to reflect later stages of somatosensory processing within the secondary somatosensory cortices [45–47]. The posterior parietal cortex also plays a role in this later processing, starting at approximately 90ms for upper limb stimulation [46,48]. Our finding of significant face-hand inhibition at ISIs of 45 and 55 ms suggests that afferent information from the face alters hand motor representations during early somatosensory processing, albeit at a relatively advanced stage of early processing. Indeed, since we observed face-hand inhibition before 70ms it likely involves S1, and despite being later than hand-hand inhibition, still reflects the phenomenon of short-latency afferent inhibition. Had it occurred at longer ISIs (closer to 100ms) we would have suggested that the inhibition reflected long-latency afferent inhibition and relied upon late somatosensory processing involves structures such as bilateral secondary somatosensory cortices and contralateral posterior parietal cortex [16,17,19,22,25].

### Arm and Forearm stimulation does not inhibit hand MEPs

The majority of afferent inhibition studies have focused on the upper and lower limbs, either looking at interactions within the same part of the limb (hand-hand [14–16,18,19,23,24,27,28] shoulder-shoulder [32], leg-leg [49], or between different limb segments (hand-arm, hand-forearm [26–28], foot-leg [29]). On the basis of these studies, it is generally believed that afferent inhibition within the upper limb is a robust phenomenon. Interestingly, however, we are only aware of one other investigation of afferent inhibition between different parts of the upper limb in which stimulation was not applied to the hand [28]. As in our study, they observed no inhibition in hand (and other) muscles following forearm stimulation at 3xSPT. Thus, it would appear that inhibition within the upper limb is not as robust as previously thought, and instead is present only when the afferent stimulation is on or near the hand.

One possible explanation for the absence of arm-hand afferent inhibition might be that the higher sensitivity and larger cortical magnification of the hand [1,2,50] leads to a larger cortical response to hand stimulation than to forearm or arm stimulation. We believe this to be unlikely, however, as a stimulus on the arm five times longer than that used in the present study still failed to inhibit hand muscle responses [28]. Given our finding of face-hand inhibition, it is important that future studies continue to investigate if and when arm-hand inhibition can be evoked.

The inhibitory interaction between the face and hand revealed here might constitute one of the sources of face-hand cortical interactions like those observed after plasticity-inducing events [3,4,9]. As initially suggested by Jacobs and Donoghue (1991), one possible mechanism of cortical reorganization is the unmasking of pre-existing lateral excitatory connections by the reduction of activity in intracortical inhibitory circuits. The inhibitory sensorimotor interaction observed here might contribute to maintaining functional boundaries between face and hand cortical territories. After a plasticity-inducing event (e.g. deafferentation), activity in the inhibitory circuitry could be decreased, resulting in the disinhibition of latent intracortical excitatory connections and a reduction in SAI, as shown by Bailey et al., (2015) in patients with spinal cord injury. Another possibility is that the face-hand sensorimotor inhibitory interactions reported here are one of the potential physiological substrates upon which a multitude of remotely represented body parts may enter a (missing) hand territory based upon the frequency of usage of these body parts [10,53,54]. Were this case, however, we should also have found AI between the arm and the hand.

In spite of increasing interest in afferent inhibition, its underlying function remains unknown (reviewed in Turco et al., 2017). Some studies use it as a tool to investigate the integrity of the cholinergic system [20,56], while others use it as we did here: as a tool to probe sensorimotor interactions in neurologically healthy individuals [15,16,22,24,25]. Participants in these studies are always seated quietly and never perform any particular task. The experiments presented in this paper constitute the first step in investigating the existence of SAI between the face and the hand as the ISIs at which it is present. In the future, it will be interesting to directly compare the amount and latency of SAI at various body sites within the same participants, as well as to examine whether the face hand interactions demonstrated here are altered as a function of the proximity of the two body parts and/or their engagement in hand-to-mouth behaviours. These types of experiments will not only shed more light on face-hand afferent inhibition, but could also help us to better understand the functional importance of the sensorimotor interactions that underlie afferent inhibition.

## Supporting information

**S1 Appendix. Peak-to-peak mean MEP amplitudes (mV)**. Excel file with peak-to-peak mean MEP amplitudes (mV) values per condition per subject used to perform the statistical analysis.

